# Structural and molecular insight into the pH-induced low-permeability of the voltage-gated potassium channel Kv1.2 through dewetting of the water cavity

**DOI:** 10.1101/775759

**Authors:** Juhwan Lee, Mooseok Kang, Sangyeol Kim, Iksoo Chang

## Abstract

Understanding the gating mechanism of ion channel proteins is key to understanding the regulation of cell signaling through these channels. Channel opening and closing are regulated by diverse environmental factors that include temperature, electrical voltage across the channel, and proton concentration. Low permeability in voltage-gated potassium ion channels (Kv) is intimately correlated with the prolonged action potential duration observed in many acidosis diseases. The Kv channels consist of voltage-sensing domains (S1–S4 helices) and central pore domains (S5–S6 helices) that include a selectivity filter and water-filled cavity. The voltage-sensing domain is responsible for the voltage-gating of Kv channels. While the low permeability of Kv channels to potassium ion is highly correlated with the cellular proton concentration, it is unclear how an intracellular acidic condition drives their closure, which may indicate an additional pH-dependent gating mechanism of the Kv family. Here, we show that two residues E327 and H418 in the proximity of the water cavity of Kv1.2 play crucial roles as a pH switch. In addition, we present a structural and molecular concept of the pH-dependent gating of Kv1.2 in atomic detail, showing that the protonation of E327 and H418 disrupts the electrostatic balance around the S6 helices, which leads to a straightening transition in the shape of their axes and causes dewetting of the water-filled cavity and closure of the channel. Our work offers a conceptual advancement to the regulation of the pH-dependent gating of various voltage-gated ion channels and their related biological functions.

**Author Summary:** The acid sensing ion channels are a biological machinery for maintaining the cell functional under the acidic or basic cellular environment. Understanding the pH-dependent gating mechanism of such channels provides the structural insight to design the molecular strategy in regulating the acidosis. Here, we studied the voltage-gated potassium ion channel Kv1.2 which senses not only the electrical voltage across the channels but also the cellular acidity. We uncovered that two key residues E327 and H418 in the pore domain of Kv1.2 channel play a role as pH-switch in that their protonation control the gating of the pore in Kv1.2 channel. It offered a molecular insight how the acidity reduces the ion permeability in voltage-gated potassium channels.

## Introduction

Electrical signals in neurons are generated by sequential gating of several voltage-gated ion channels on their cell membranes. The opening and closing of these channels are not only sensitively controlled by membrane potentials in general, but also respond to the intra- and extracellular conditions, such as chemicals [1, 2], mechanical pressure [3], temperature [4], and proton concentrations [5]. Among these channels, the voltage-gated potassium channels (Kv) are selectively permeable to potassium ions and repolarize the membrane potential in response to depolarizing voltage [6]. The molecular mechanisms underlying this potassium ion-selectivity and voltage-dependent gating of the Kv channels have been extensively studied [7-11]. However, the molecular mechanism of pH-dependent gating in Kv channels is less well-understood, although it has been revealed that the potassium ion permeability is inhibited by the high proton concentration in acidosis [12]. The low permeability in the Kv channels is intimately correlated with the prolonged action potential duration observed in acidosis diseases such as cardiac arrhythmias.

These Kv channels have a tetrameric structure composed of four homo-subunits surrounding an ion-transporting pore, with each subunit containing six membrane-spanning α-helices called S1–S6. They are spatially separated from a voltage-sensing domain-containing S1–S4 and a central pore domain-containing S5–S6 that includes a P-helix (Fig 1A). These two domains are connected by an S4–S5 helical linker [13]. The pore domain contains a potassium ion-selective pathway and gates spanning the cell membrane. The narrowest part (extracellular side) on the pore in the channels is the “selectivity filter”, whereas the opposite part (intracellular side) of the filter on the pore is the “water-filled cavity” (Fig 1B). The gating of the Kv channels is structurally determined by whether the water-filled cavity is wetted or dewetted.

**Figure 1.**
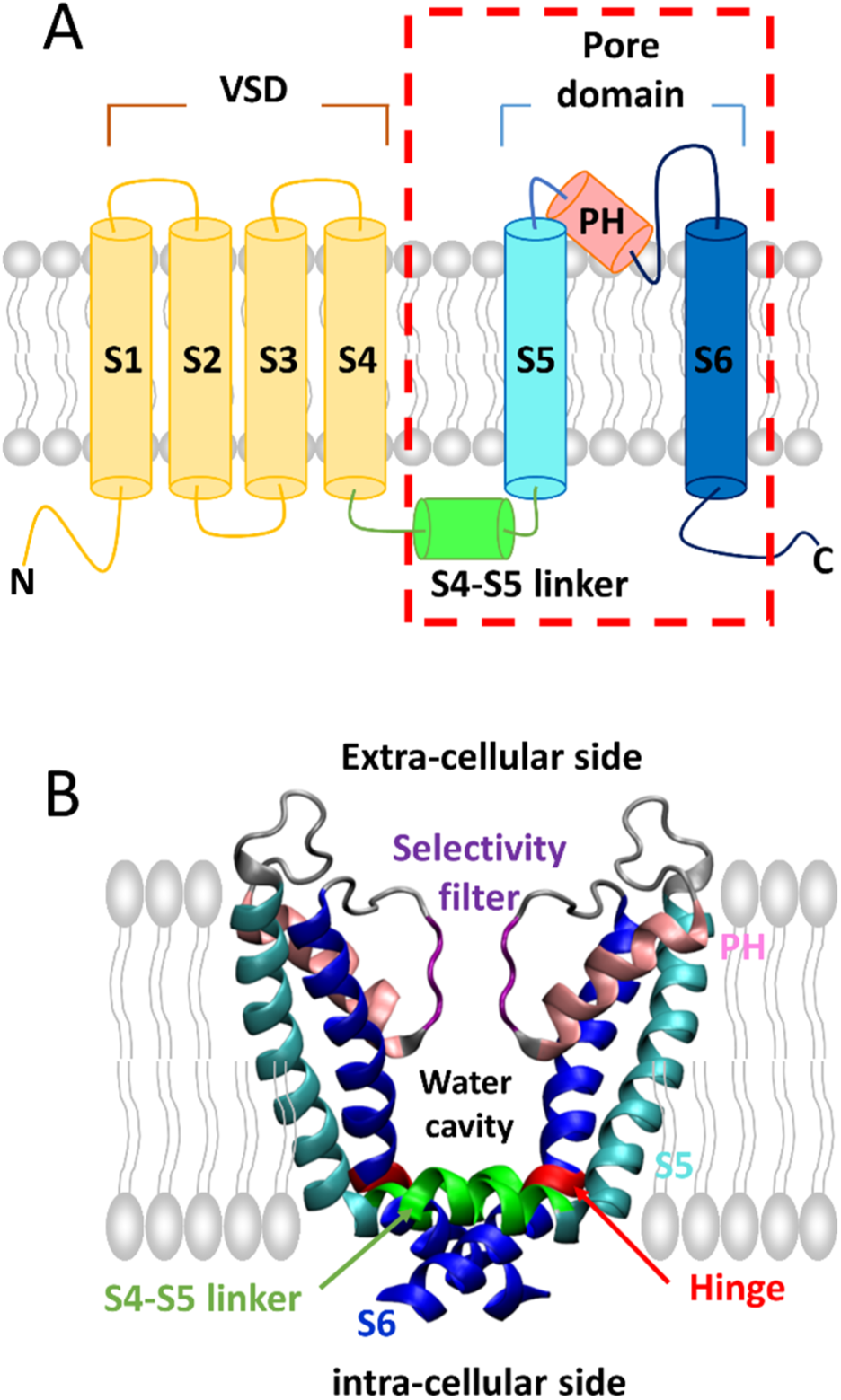
Schematic illustration of the Kv channel structure. **(A)** A cartoon model of a subunit of the Kv channel showing the voltage-sensing domains (VSDs) (S1–S4) in yellow, the S4–S5 helical linker in green, the S5 helix in cyan, the P-helix in pink, and the S6 helix in blue. **(B)** Two opposite subunits of the pore domain of the Kv channel are represented by ribbons. The other two subunits have been omitted. The selectivity filter is shown in purple, and the water cavity is located below the water cavity. The hinge region in S6 helix is highlighted in red.

In Kv1.2 channels, as in the other members of the non-inactivating ion channel family, ionic currents flow in response to the applied depolarized voltage and are maintained until the end of depolarization. Kv1.2 channels maintain the closed conformation at polarized (resting) potentials or the open conformation at depolarized potentials. The ionic currents are intimately connected to the permeability of the ions in the channels. Previous experimental work showed that the permeability of Kv1.2 channels to potassium ions gradually decreases as the pH changes from 7.5 to 4.5, with the experimental permeability midpoint occurring at around pH 5.3 [12]. This implies that the protonation of some titratable residues make the channels resist the transition from closed to open conformations against depolarized voltage.

Here, we report our observations from an all-atom molecular dynamics (MD) simulation. We found that the voltage-gated Kv1.2 ion channel has dual-functionality due to protonation of the conserved residues E327 and H418 situated near the water cavity on the intracellular side because it induces the gating transition of the pore domain from an open to closed position under acidic conditions. We characterized the structural role of the two determinant residues E327 and H418 in the gating of Kv1.2 channels, which can be protonated under a reasonable acidic condition, as demonstrated in the previous experimental study [12]. We suggest the molecular and structural mechanism underlying the acid-induced low permeability of Kv1.2 channels by uncovering the mechanism through which the channels, under acidic conditions, prefer the closed-pore state with the dewetted cavity than the open-pore state with the wetted cavity.

## Results

### Identification of the key acid-responding residues in Kv1.2

To identify the key residues that might be protonated under acidic conditions, that is, from pH 7.5 to around pH 5.3, we attempted to estimate the pKa values of the titratable residues in Kv1.2 channels with closed conformations. Because only the open conformations of the Kv1.2 channel were available from the Protein Data Bank [14], we reconstructed the closed conformations of the Kv1.2 channels through MD simulations, starting with the initially open conformation after protonation of three residues (E327, H418, and E420) in the proximity of the water-filled cavity (Fig S1). Here, we considered the pore domains (residue numbers: 312– 421) that are responsible for the gating of the Kv1.2 channel, excluding the voltage-sensing domains. Based on the structural ensembles of the closed conformations, the pKa values of the titratable residues in the channel were estimated (Fig 2A) [15]. The pKa values of both E327 in the A- and D-chain and H418 were around 6.0, which means that these two residues can be protonated under the above acidic conditions. Multiple sequence alignment for Kv1 subfamily proteins showed that these two residues are conserved (Fig 2B) and are located near the water-filled cavity (Fig 2C). Therefore, we decided to investigate the effect of protonation of the E327 and H418 residues on the conformational transition of the pore domain of the Kv1.2 channel.

**Figure 2.**
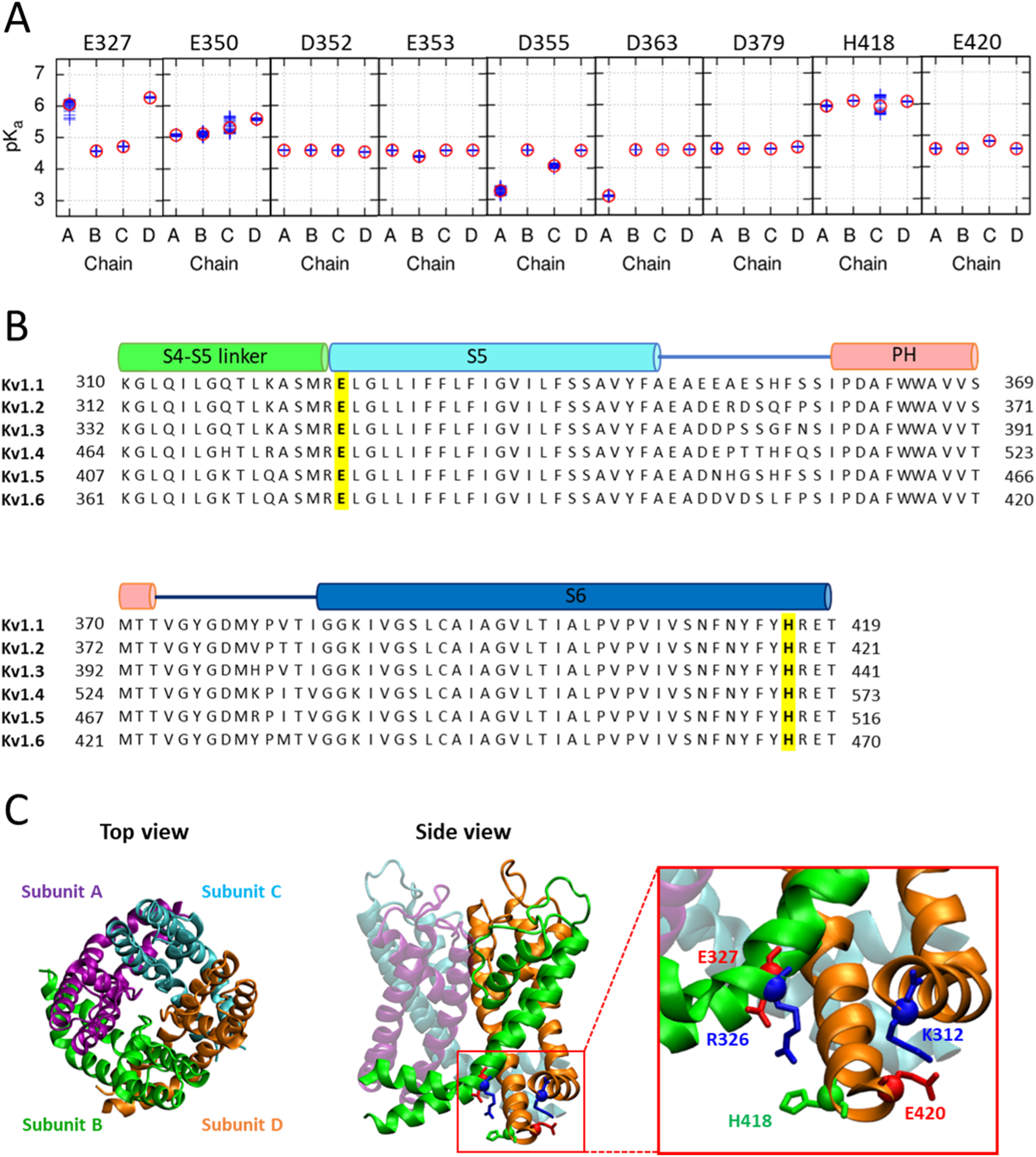
pKa estimates of H418 and acidic residues and two key residues in the Kv1.2 channel. **(A)** The pKa estimates of H418 and the acidic residues in the pore domain of the closed Kv1.2 channels. **(B)** Multiple sequence alignment for the pore domains of the rat Kv1 subfamily. The cylinders above the sequences denote the secondary structural information of the four helical segments. The two key conserved residues E327 and H418 are highlighted in yellow and bold font. **(C)** Top and side views of the open pore domain of the Kv1.2 channel. The four subunits are represented by different colors in the ribbon diagram. The magnified view shows the key residues E327 and H418 and their neighbors in the inner helical bundle, which are represented by the licorice and Cα balls.

### Protonation of E327 and H418 induces closure of Kv1.2

We performed atomistic MD simulations of the central pore domain in the Kv1.2 channel for both a Ep327/Hp418 state and a wild-type (as a control group) (here, “p” indicates the protonation; details of our MD simulations are provided in the Supplementary Information). Starting from the initial open conformation of the central pore domain in the Kv1.2 channel, five trajectories of MD simulations for each of a wild-type and a Ep327/Hp418 state were run for 2 μs, and the equilibrium configurations of the central pore domain in the Kv1.2 channel were obtained (Fig 3A). The time evolution of the number of water molecules in the cavity of the pore showed that the conformations from three trajectories of the Ep327/Hp418 state equilibrated to the closed conformations with the dewetted cavity, whereas those from all five trajectories of the wild-type remained in the open conformations with the wetted cavity (Figs 3B and 3C) [16]. This result demonstrates that the pore domain of Kv1.2 channels prefers the closed conformations under acidic conditions, whereas the open conformations are abundant under the neutral pH condition. Therefore, acidic conditions hamper the opening of the channel even under a depolarizing voltage and lower the ion permeability of the channel.

**Figure 3.**
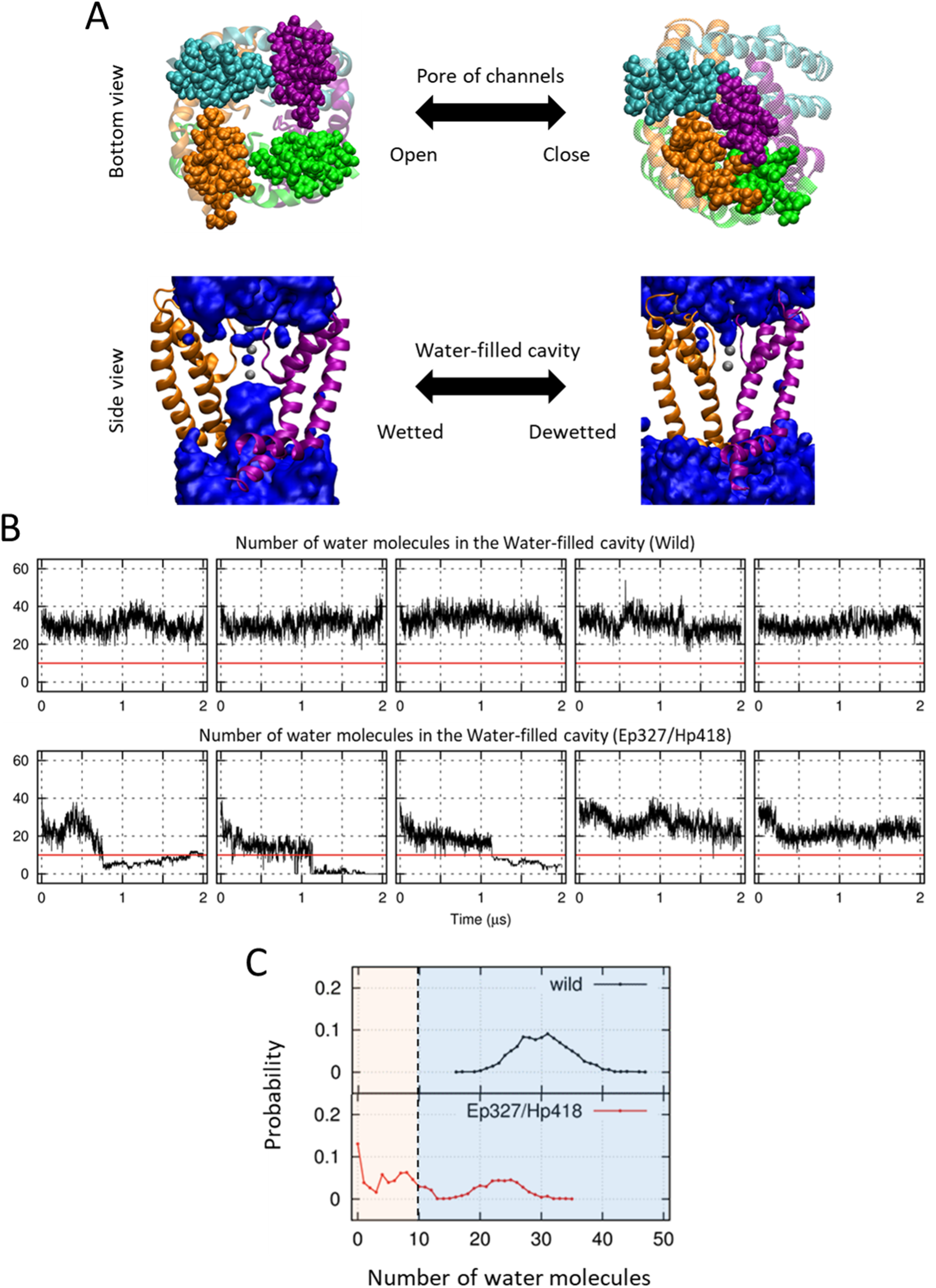
Pore closure due to protonation of the two key residues E327 and H418 in the Kv1.2 channel. **(A)** The structures of the open and closed Kv1.2 channel. Here, the ribbons represent the pore domain of Kv1.2 channels, the blue spheres denote water molecules, and the gray spheres denote potassium ions. The upper panel illustrates the bottom view of the open and closed conformations. The lower panel shows the wetted and dewetted water cavity in the channel. Both the two opposite subunits and lipid molecules have been omitted from the lower panel. **(B)** The time evolution of the number of water molecules in the water cavity of the wild-type or Ep327/Hp418 states of the Kv1.2 channel. The five individual plots in each state represent the simulation results from each trajectory of our MD simulations. In addition, the red horizontal line separates the wetted state from the dewetted state of the water cavity. **(C)** The distributions of the number of water molecules in the water cavity. The left region with fewer water molecules corresponds to the dewetting condition, whereas the right region with many water molecules corresponds to the wetting condition.

### Structural and molecular mechanisms of the pore closure of Kv1.2

Based on the structural ensembles collected from our MD simulations, we quantified the structural alterations between the wild-type and Ep327/Hp418 type of Kv1.2 channels. The hinge of the S6 helix maintains electrostatic balance through two inter-subunit interactions of R326–H418 and E327–H418. These interactions stabilize the open conformation of the Kv1.2 pore domain under a neutral pH condition (left panel of Fig 4A). We used two structural determinants that can distinguish the open conformation from the closed conformation of the pore in Kv1.2 channels, namely, the R326–H418 inter-subunit distance defined by the nearest inter-atom distances in these two residues (the bottom view in Fig 4A) and the dihedral angle extended by the positions of the Cα atoms in four residues (R326, L400, V408, and Y415) on the S6 helix (the side view in Fig 4A). The dihedral angle determines whether the S6 helix is bent or straight [17].

**Figure 4.**
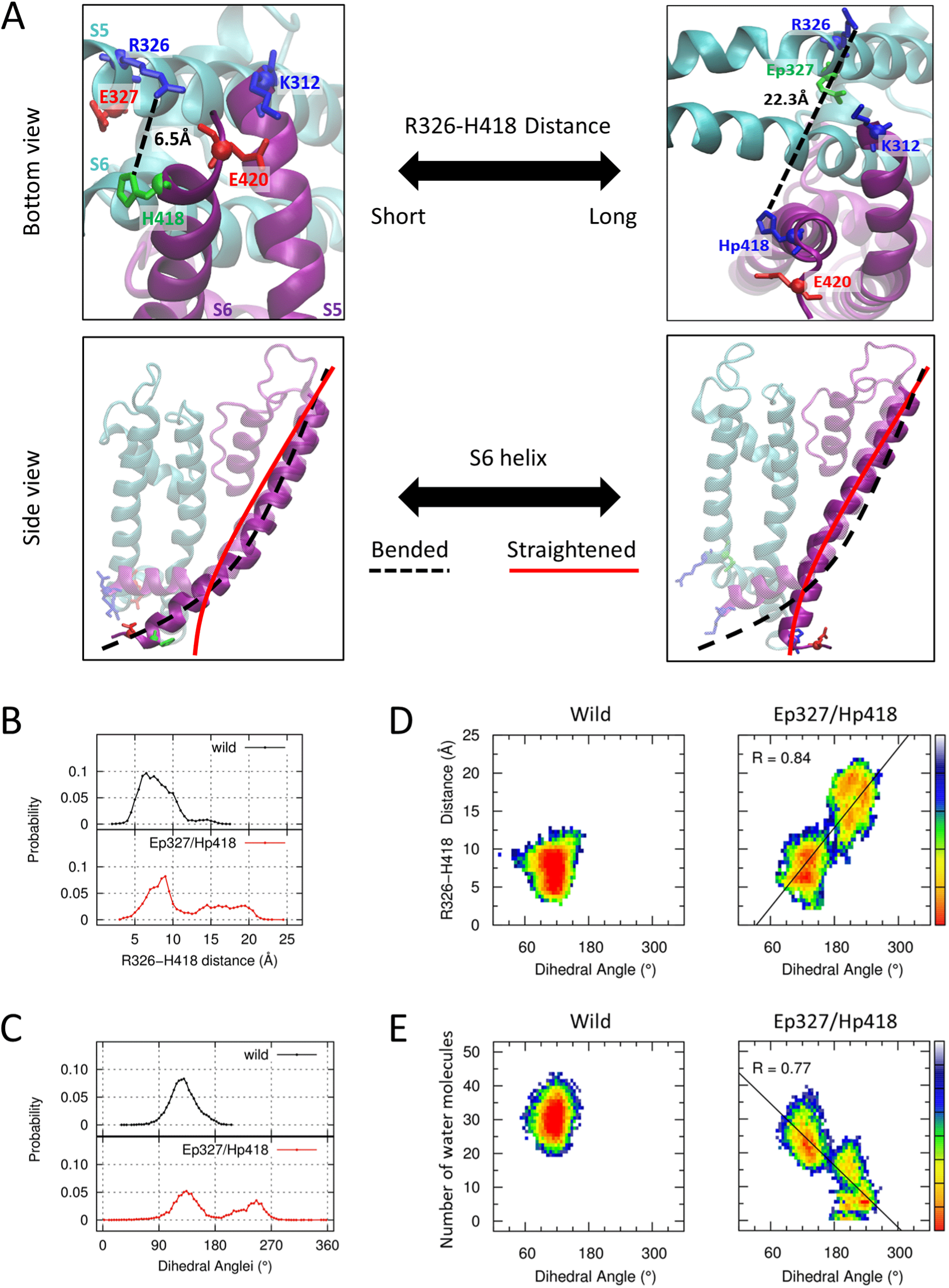
Straightening of the S6 helix due to protonation of the two key residues E327 and H418 in the Kv1.2 channel. (**A)** The detailed structures of the open (left panels) and closed (right panels) Kv1.2 channels. The upper panel illustrates the distance between the R326 and H418 residues. For the open conformation of the pore, the distance between E327 and H418 is 7.6 Å, whereas it is 14.7 Å for the closed conformation. In addition, the lower panel shows the S6 helix when it is bent (black dotted line) or straight (solid red line). Here, two neighbor subunits are displayed using purple and cyan color, respectively, while the other subunits have been omitted. **(B)** The probability distribution curves for the inter-subunit distances between the R326 and H418 residues of the wild-type and Ep327/Hp418 states. **(C)** The probability distribution of the dihedral angles, extended by the position of the Cα atoms in R326, L400, V408, and Y415, for each state. The left half region with the angle smaller than 180° corresponds to the bent S6 helix, whereas the right half and the secondary peak is the straight S6 helix. **(D)** Free energy landscapes (in arbitrary units) for the R326–H418 distances and the dihedral angles. In the right panel for the Ep327/Hp418 state, a high correlation value of 0.84 was detected. **(E)** Free energy landscapes (in arbitrary units) for the dihedral angles and the number of water molecules in the water-filled cavity. In the right panel for the Ep327/Hp418 state, a high correlation value of 0.77 was also detected.

The Supplementary Information in Figs S2 and S3 shows the time evolution of the values of the R326–H418 inter-subunit distance and the dihedral angle from the structural ensembles collected in the last 500 ns time window of our MD simulations, which demonstrates the effects of the protonation of E327 and H418. The R326–H418 distances in the Ep327/Hp418 state of the Kv1.2 channels become longer and the distance distribution is much broader compared with those of the wild-type, for which the most frequent distances are around 6 Å (Fig 4B). The drastic increase in the R326–H418 distances in the Ep327/Hp418 state is due to the repulsive Coulomb interaction between R326 and Hp418. The distribution of the dihedral angles extended by R326, L400, V408, and Y415 on the S6 helix of the wild-type Kv1.2 channels has a distinct peak at around 130°, indicating that the S6 helix is bent (the dotted black line in Fig 4C). On the other hand, the protonation of both E327 and H418 gives rise to a secondary peak around 245° in their distribution, indicating that the S6 helix is straightened (the solid red line in Fig 4C). The increase in the R326–H418 distance is closely correlated with the increase in the dihedral angle extended by R326, L400, V408, and Y415 on the S6 helix and is well captured by the heat map of the ensemble population along the two axes of each quantity (Fig 4D). The heat map in the Ep327/Hp418 state revealed that the increase in the R326–H418 inter-subunit distance straightened the S6 helix. The change in the degree of the dewetting in the cavity of the Kv1.2 channels is also closely correlated with the straightening of the S6 helix. This close correlation is demonstrated in the heat map of the ensemble population along the two axes of the dihedral angle and in the number of water molecules in the cavity (Fig 4E).

Overall, Hp418 was electrostatically pushed by R326 at the hinge of the S6 helix under acidic conditions. This changed the shape of the S6 helix from bent to straight. Therefore, the straightening of the S6 helix is a robust indication of pore closure in the potassium channel Kv1.2 [18-20]. Here, we suggest that the repulsive Coulomb interaction of the inter-subunits triggered by the protonation of the two key residues E327 and H418 is the molecular mechanism of the pore closure in the Kv1.2 channels, together with both the increase in the inter-subunit distance and the straightening of the S6 helix.

### Altering the permeability of Kv1.2 through mutation of E327 and H418

The permeability of the Kv1.2 channel was further probed by examining the changes in the inter-subunit interactions among R326, E327, and H418 and the intra-subunit interaction between K312 and E420. We were able to modify the charge states of these residues through protonation or mutation, as shown in Table 1. First, we modified the inter-subunit interaction by changing the charge state of H418 to resemble the weak acidic condition. Second, we changed the charge states of both E327 and H418 to break the inter-subunit interaction, which resembled the stronger acidic condition. Third, we modified the intra-subunit interaction by changing the charge state of E420. Finally, we broke both the inter- and intra-subunit interactions by simultaneously changing the charge state of E327, H418, and E420, which resembled the strong acidic condition. Our simulation results showed that the transition from the open to the closed conformation in the pore of the Kv1.2 channel depends on the state of the inter-subunit interaction (Table 1). The E327 and H418 residues located on the intracellular side across the membrane have a major effect on the pH-sensing mechanism, depending on their protonation states.

**Table 1.**
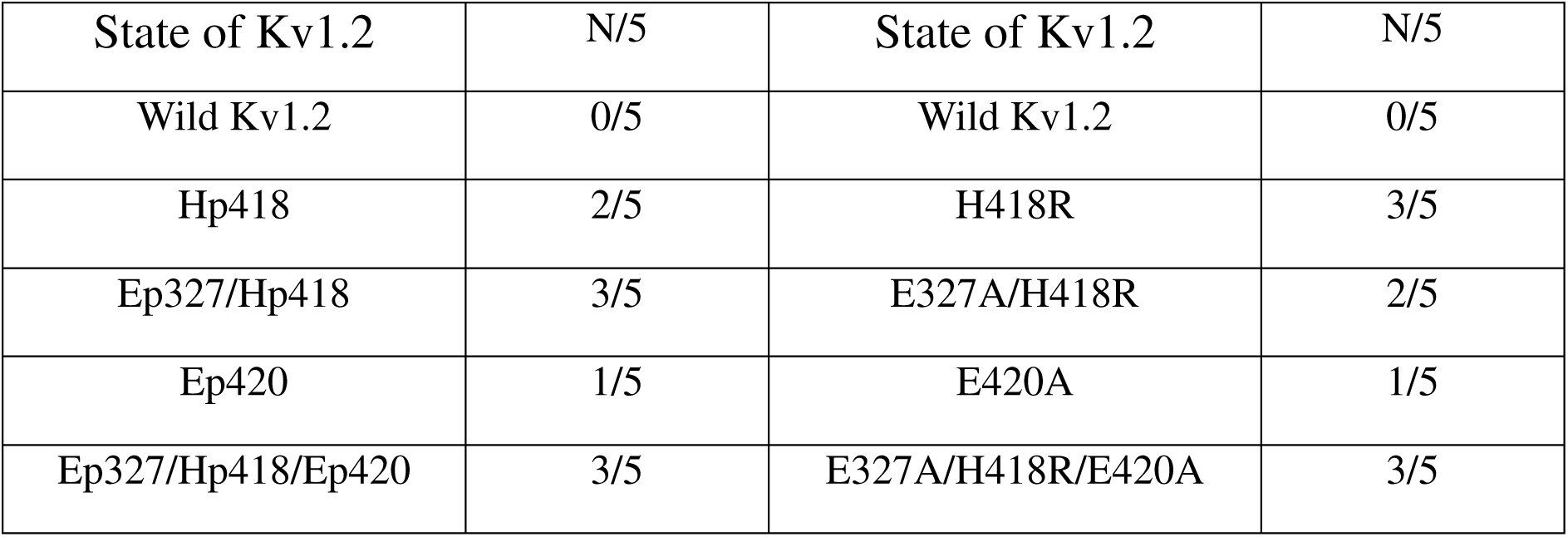
List of mutants subjected to MD simulation and the number of trajectories equilibrated to the closed conformations with the dewetted cavity out of five trajectories (N/5).

E327/H418R mutant mimics the effect of acidic pH conditions. As expected, our simulation results showed that the pore domain of the mutant prefers the closed conformations compared with the wild-type Kv1.2 channel (Table 1). In addition, the molecular mechanism of the pore closure of the mutant was similar to that of the Ep327/Hp418 state of Kv1.2, showing both an increase in the inter-subunit distance and a straightening of the S6 helix (right panels of Figs 5A and 5B). For the pKa estimation of each titratable acidic residue in each subunit of the Kv1.2 channel, all four H418 residues in the four subunits were protonated under acidic pH conditions, whereas only two E327 residues in the four subunits were protonated (Fig 2A). Thus, we believe that the H418 residue is a more suitable target of a single mutation than E327. As expected, our simulated H418R mutant preferred the closed conformations compared with the wild-type Kv1.2 channel (Table 1). In the left panels of Figs 5A and 5B, the closed conformation of the H418R mutant almost resembled the closed conformations of the Ep327/Hp418 state and the E327A/H418R mutant. Based on the molecular mechanism of the Kv1.2 channel, we were thus able to control the pore closure of Kv1.2 through mutation of the E327 and/or H418 residues.

**Figure 5.**
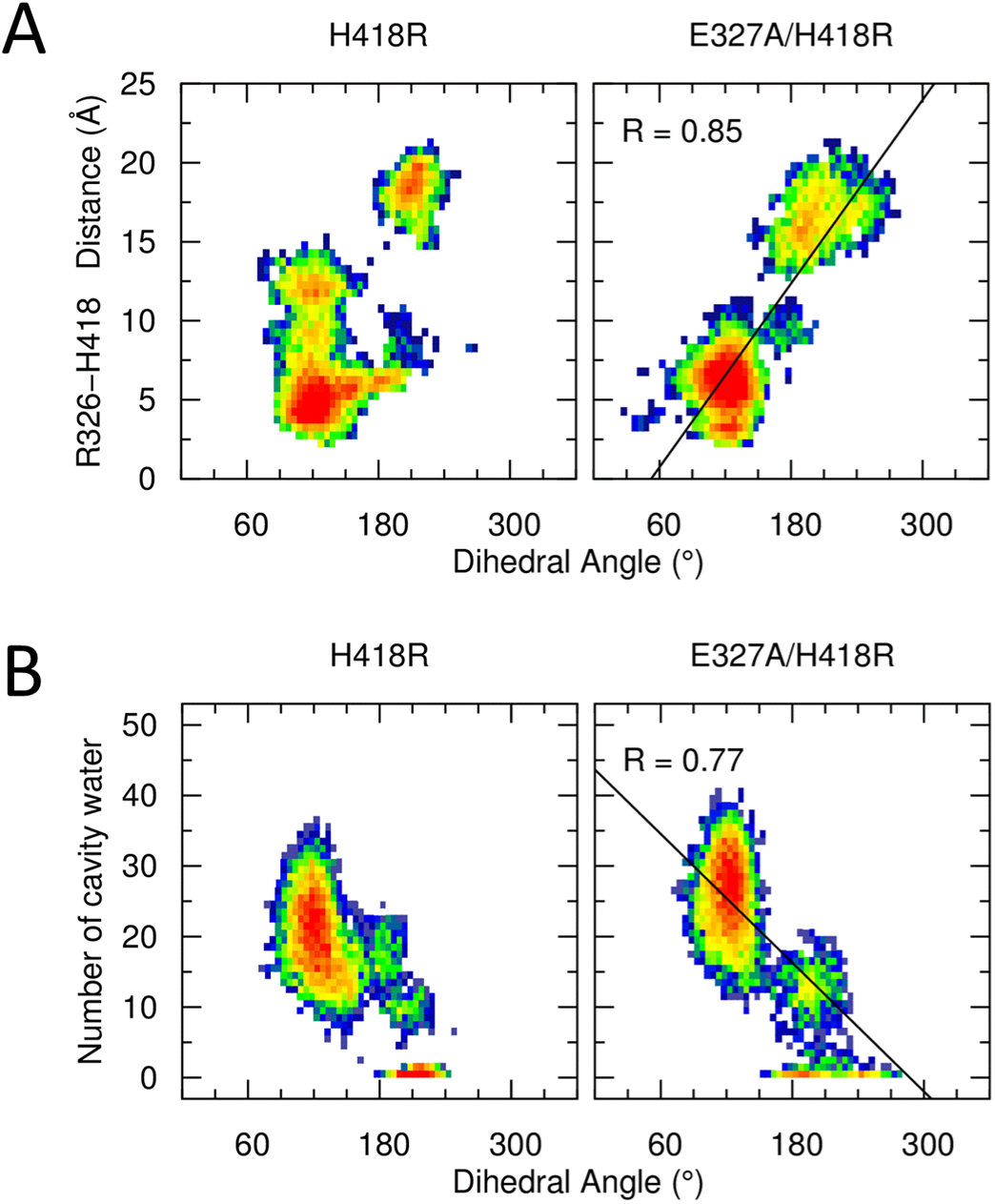
Descriptions of the effects of H418R and E327A/H418R mutants. **(A)** Free energy landscapes (in arbitrary units) for the R326–H418 distances and dihedral angles. In the right panel for the E327A/H418R state, a high correlation value of 0.85 was detected. **(B)** Free energy landscapes (in arbitrary units) for the dihedral angles and number of water molecules in the water-filled cavity. In the right panel for the E327A/H418R state, a high correlation value of 0.77 was also detected.

## Discussion

We investigated the molecular mechanism underlying the pH-dependent gating of the pore domain of the Kv1.2 channel protein under intracellular acidic conditions. A decrease in environmental pH from 6 to 5 causes Kv1.2 to undergo a conformational change from open to closed [12]. Our pKa estimates indicate that only two amino acids E327 and H418 change their charge states in response to a change in the environmental pH. Thus, E327 and H418 are proposed as key residues for pH sensing. To assess the role of the key residues under acidic pH conditions, we performed MD simulations with key protonated residues. Ep327 and Hp418 highly destabilize the electrostatic interaction. The inter-subunit interaction, which consists of R326, E327, and H418, is critically destabilized by the change in the charge states of glutamic acid and histidine. Because the net charge of the inter-subunit network is changed from 0 to +2, the repulsive force between R326 and Hp418 causes the distance to increase from 6.5 Å to 22.3 Å. This repulsive force pushes the end of S6, leading to its distortion. In our simulation, the channels are perfectly closed when the two opposite subunits of S6 undergo a conformational change. Two distorted S6 helices move close together and fill the space previously occupied by the water. As a result, the low permeability of Kv1.2 under acidic conditions was induced by the protonated E327 and H418. We tracked the step-by-step conformational change using MD simulation. Additionally, our mutant studies showed that the inter-subunit interaction (R326, E327, and H418) is more important than the intra-subunit interaction (K312 and E420). Channel gating is controlled by the charge states of the inter-subunit interaction rather than the intra-subunit interaction. This MD simulation result indicating that E327 and H418 are important for channel gating is consistent with our pKa estimate. H418R and E327A/H418R mutants undergo a structural change from an open to closed conformation. In our simulation, H418R and E327A/H418R mutants have low ion permeability, even under the neutral pH condition, and we were able to genetically modulate the ion permeability of Kv1.2 via mutation of the key residues.

The molecular mechanism of the voltage-dependent gating of the Kv channels involves displacement of the voltage-sensing domains that regulate the wetting or dewetting of the cavity [7]. Therefore, the coexistence of these voltage- and acid-dependent gating mechanisms in Kv1.2 channels implies that both the voltage-induced structural pressure and acid-induced structural pressure can simultaneously influence the gating of the channels. The possibility for this simultaneous action of two different mechanisms of pore gating was indicated in previous experimental studies of the behavior of the potassium ion current of Kv1.2 channels that suggested that the acidity competes with the depolarizing voltage [12]. At a glance, this might be counterintuitive because the neuronal channels were not able to distinguish the environmental factor that played a role in their own gating. However, it is worth noting that the time scale for the fluctuation in the membrane potential is on the order of milliseconds, whereas that of the cellular pH is from seconds to minutes. Thus, the permeability of potassium ion currents through Kv1.2 channels is not only finely controlled by the depolarizing voltage in the short time interval but also governed by the acidity in the long time interval. Our study offers insight into the dual-gating mechanisms of the Kv channels, which orchestrate both the voltage-dependent and pH-dependent gating mechanisms of different molecular mechanisms.

## Methods

### All-atom molecular dynamics simulation

We performed MD simulations using GROMACS 5.0 [21] with the CHARMM36 force field [22]. The initial configurations of the Kv1.2 channel for the MD simulation were generated using CHARMM-GUI lipid builder [23]. The system consists of a channel protein, lipids (271 POPE in upper and lower leaflets), water molecules (∼ 10,000 TIP3P water molecules), and ions (0.6 M; K^+^, 133; Cl^−^, 125; wild-type condition). The channel protein is a symmetric tetramer structure with an open-pore domain (residues 312–421) that is derived from the X-ray crystal structure of the Kv1.2 channel (PDB code: 2R9R) [14]. The system includes ∼ 71,000 atoms in a rectangular box (100 × 100 × 75 Å) under the periodic boundary condition. The particle-mesh Ewald (PME) method [24] was applied for assessing long-range electrostatic interactions with a 12 Å cut-off distance, and potential-based switching functions were used for van der Waals interactions with a 10–12 Å switching range. The position of the hydrogen atoms was restrained by the equilibrium bond length using the LINCS algorithm [25]. Approximately 5,000 steps of steepest-descent minimization and 20,000 steps of equilibrating simulation were conducted. An additional 30 ns simulation with a restrained protein backbone was performed to equilibrate the water cavity position, followed by a production run. The production simulations were carried out for 2 μs with a 2 fs time step in NPT ensemble holding a constant particle number (N = ∼ 71,000), pressure (P = 1 bar), and temperature (T = 310 K). Temperature was controlled by the Nosé–Hoover temperature coupling method [26, 27] with a tau-t of 1 ps and pressure was maintained by the semi-isotropic Parrinello–Rahman method [28, 29] with a tau-p of 5 ps and compressibility of 4.5 × 10^−5^ bar^−1^. All trajectories were recorded every 10 ps, and VMD software [30] was used for the visual analysis.

### pKa calculation

For pKa estimates of H418 and the acidic residues in the pore domain of the closed Kv1.2 channels, we extracted 98 ensemble structures in the last 1-μs time window from our MD simulation for a Ep327/Hp418/Ep420 state. In addition, after applying MEAD programs [15], we selected 765 titration states of the channels that were predominant in the pH range from 3 to 8. At each titration state, we calculated the energy for each of 98 ensemble structures, as well as their average and standard deviation using AMBER force field 99SB [31]. Red circles in the figure represent the pKa values when the average energies of ensemble structures, ⟨*E*⟩, were used to calculate the titration fraction as a function of pH for the corresponding titration states. The blue crosses represent the pK_a_ values when the statistical variance of the energies ⟨*E*⟩ ± 0.1σ, ⟨*E*⟩ ± 0.2σ, …, ⟨*E*⟩ ± 1σ was considered to reflect the flexibility of the pK_a_ values due to the structural fluctuation of ensemble structures.

## Acknowledgments

We acknowledge the DGIST Supercomputing Bigdata Center for the allocation of dedicated supercomputing time. This study was supported by the Creative Research Initiatives of the National Research Foundation (NRF) of Korea (2008-0061984) and the DGIST Core Protein Resources Center funded by MOTIE, Korea (N0001822).

## Supporting Information

**S1 Fig. Time evolution of the number of water molecules in the water cavity of the Ep327/Hp418/Ep420 states of the Kv1.2 channel, related to Figure 2.** Starting from the initially open conformation of the Kv1.2 channel with the wetted cavity, we protonated E327/H418/E420 and conducted our MD simulations. The MD simulation trajectory after 550 ns shows that the number of water molecule in the cavity drops to near zero, demonstrating that the initially wetted cavity becomes dewetted. This result implies that the conformation of the pore of the Kv1.2 channel changed from the open conformation to the closed conformation.

**S2 Fig. Time evolution curves of the distance between the R326 and H418 residues from five individual trajectories for the wild-type and Ep327/Hp418 states of the Kv1.2 channel, related to Figure 4.** The four subunits of the pore domain of the Kv1.2 channel are denoted using different colors (purple, green, cyan, and yellow).

**S3 Fig. Time evolution curves of the dihedral angle from five individual trajectories for the wild-type and Ep327/Hp418 states of the Kv1.2 channel, related to Figure 4.** The four subunits of the pore domain of the Kv1.2 channel are denoted using different colors (purple, green, cyan, and yellow).

## Supplementary Material

### Supplemental Figures

**Figure S1.**
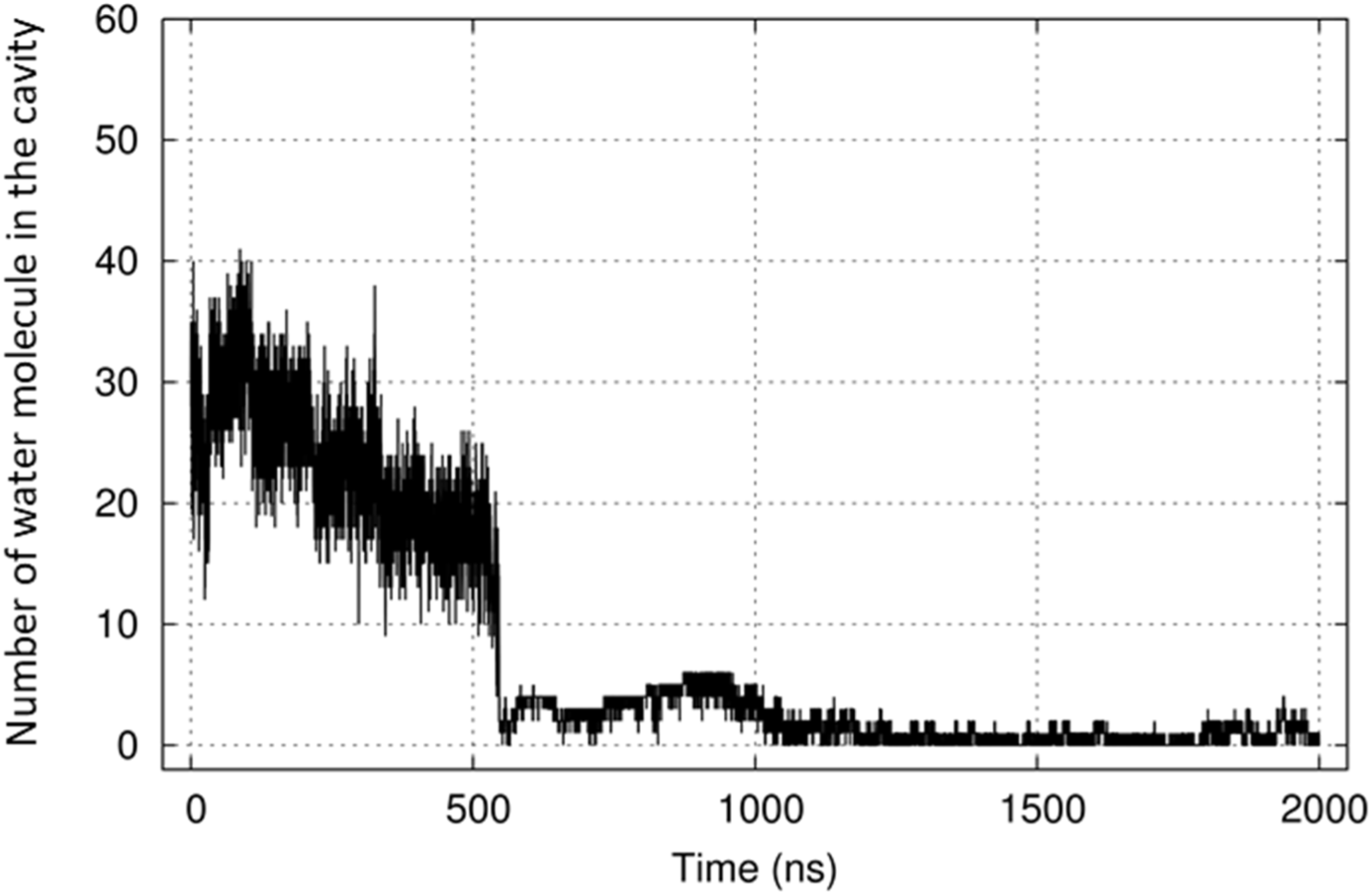
Time evolution of the number of water molecules in the water cavity of the Ep327/Hp418/Ep420 states of the Kv1.2 channel, related to Figure 2. Starting from the initially open conformation of the Kv1.2 channel with the wetted cavity, we protonated E327/H418/E420 and conducted our MD simulations. The MD simulation trajectory after 550 ns shows that the number of water molecule in the cavity drops to near zero, demonstrating that the initially wetted cavity becomes dewetted. This result implies that the conformation of the pore of the Kv1.2 channel changed from the open conformation to the closed conformation.

**Figure S2.**
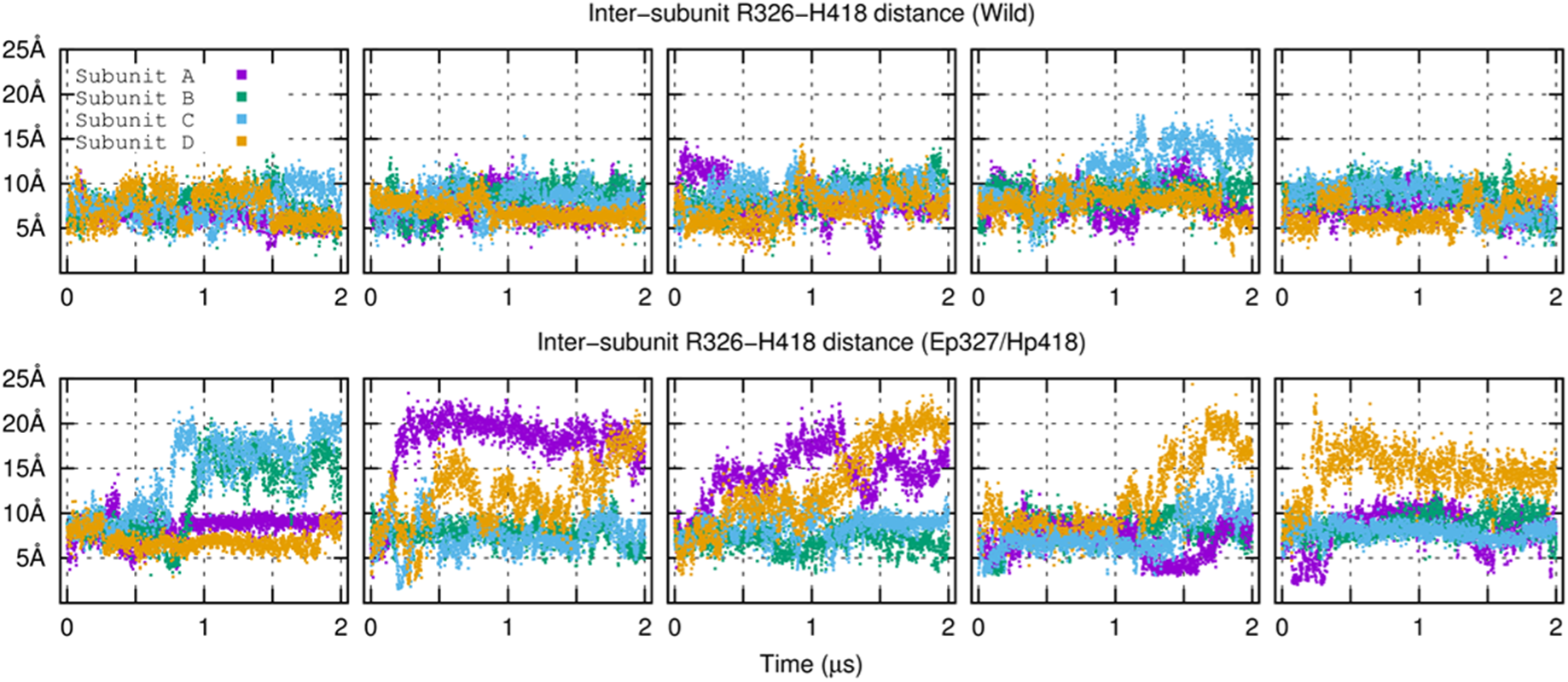
Time evolution curves of the distance between the R326 and H418 residues from five individual trajectories for the wild-type and Ep327/Hp418 states of the Kv1.2 channel, related to Figure 4. The four subunits of the pore domain of the Kv1.2 channel are denoted using different colors (purple, green, cyan, and yellow).

**Figure S3.**
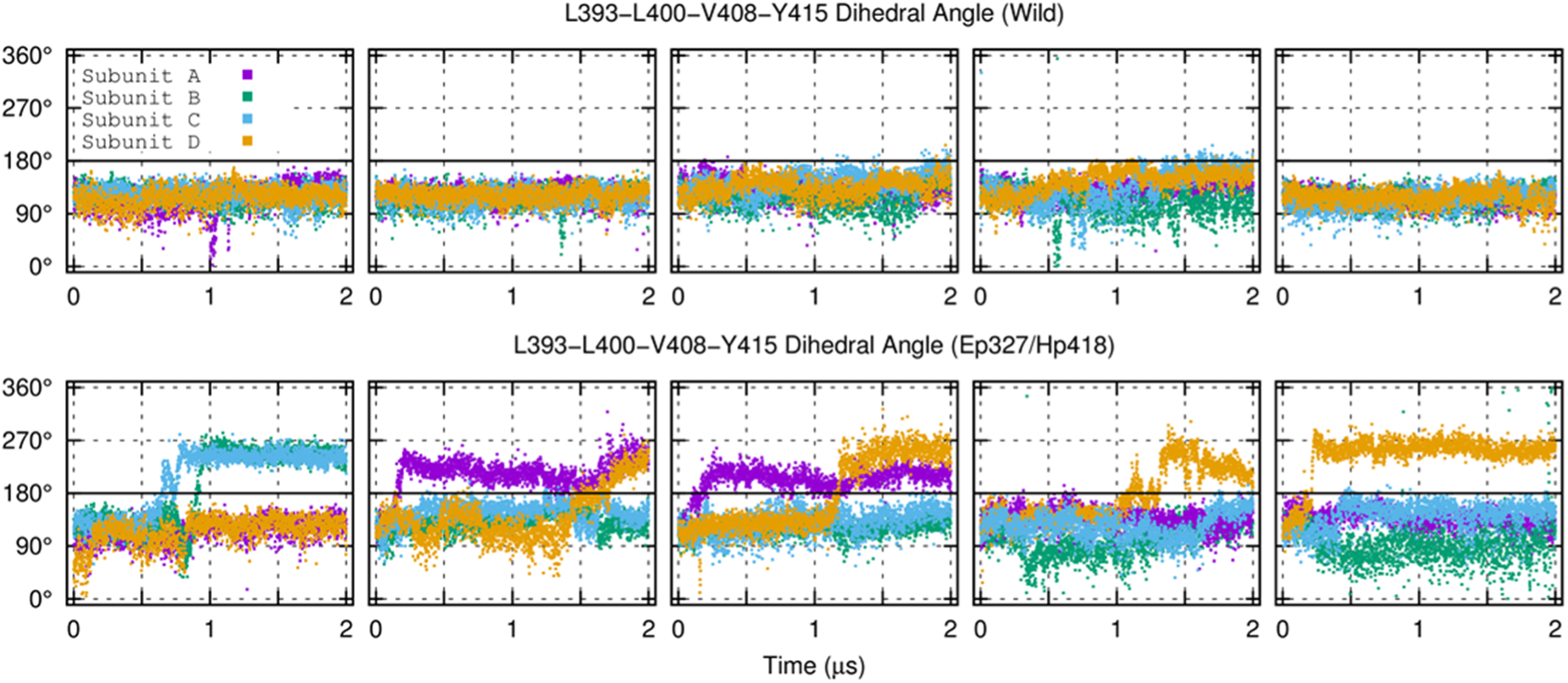
Time evolution curves of the dihedral angle from five individual trajectories for the wild-type and Ep327/Hp418 states of the Kv1.2 channel, related to Figure 4. The four subunits of the pore domain of the Kv1.2 channel are denoted using different colors (purple, green, cyan, and yellow).

